# Potential consequences of the red blood cell storage lesion on cardiac electrophysiology

**DOI:** 10.1101/2020.05.22.111302

**Authors:** Marissa Reilly, Chantal Bruno, Tomas Prudencio, Nina Ciccarelli, Devon Guerrelli, Raj Nair, Manelle Ramadan, Naomi L.C. Luban, Nikki Gillum Posnack

**Affiliations:** Sheikh Zayed Institute for Pediatric Surgical Innovation, Children’s National Hospital, Washington DC USA 20010; Children’s National Heart Institute, Children’s National Hospital, Washington DC USA 20010; Division of Critical Care Medicine, Children’s National Hospital, Washington DC USA 20010; Division of Hematology and Laboratory Medicine, Children’s National Hospital, Washington DC USA 20010; Department of Pediatrics, George Washington University, School of Medicine, Washington DC USA 20037; Department of Pathology, George Washington University, School of Medicine, Washington DC USA 20037; Department of Pharmacology & Physiology, George Washington University, School of Medicine, Washington DC USA 20037

**Author notes:** Corresponding author: Nikki Gillum Posnack, Ph.D., Sheikh Zayed Institute, 6^th^ floor, M7707, 111 Michigan Avenue, NW, Washington, DC, USA 20010. Authors contributed equally (Marissa Reilly, Chantal Bruno). AUTHOR CONTRIBUTIONS: MR^1^, NC, TP, CB, DG, MR^2^, RN performed experiments; MR^1^, NC, TP, DG and NGP analyzed data; MR^1^, NC, TP, DG and NGP prepared figures; MR^1^, TP, NC, CB, NL and NGP drafted manuscript; NL and NGP conceived and designed experiments; MR^1^, CB, NC, TP, DG, MR2, RN, NL and NGP approved manuscript. ^1^Marissa Reilly, ^2^Manelle Ramadan.

**Keywords:** red cell storage lesion, cardiac electrophysiology, hyperkalemia

## Abstract

The red blood cell (RBC) storage lesion is a series of morphological, functional and metabolic changes that RBCs undergo following collection, processing and refrigerated storage for clinical use. Since the biochemical attributes of the RBC unit shifts with time, transfusion of older blood products may contribute to cardiac complications, including hyperkalemia and cardiac arrest. We measured the direct effect of storage age on cardiac electrophysiology and compared with hyperkalemia, a prominent biomarker of storage lesion severity. Donor RBCs were processed using standard blood banking techniques. The supernatant was collected from RBC units (sRBC), 7-50 days post-donor collection, for evaluation using Langendorff-heart preparations (rat) or human stem-cell derived cardiomyocytes. Cardiac parameters remained stable following exposure to ‘fresh’ sRBC (day 7: 5.9±0.2 mM K^+^), but older blood products (day 40: 9.7±0.4 mM K^+^) caused bradycardia (baseline: 279±5 vs day 40: 216±18 BPM), delayed sinus node recovery (baseline: 243±8 vs day 40: 354±23 msec), and increased the effective refractory period of the atrioventricular node (baseline: 77±2 vs day 40: 93±7 msec) and ventricle (baseline: 50±3 vs day 40: 98±10 msec) in perfused hearts. Beating rate was also slowed in human cardiomyocytes after exposure to older sRBC (−75±9%, day 40 vs control). Similar effects on automaticity and electrical conduction were observed with hyperkalemia (10-12 mM K^+^). This is the first study to demonstrate that ‘older’ blood products directly impact cardiac electrophysiology, using experimental models. These effects are likely due to biochemical alterations in the sRBC that occur over time, including, but not limited to hyperkalemia. Patients receiving large volume and/or rapid transfusions may be sensitive to these effects.

**New & noteworthy:** We demonstrate that red blood cell storage duration time can have downstream effects on cardiac electrophysiology, likely due to biochemical alterations in the blood product. Hyperkalemia and cardiac arrest have been reported following blood transfusions, but this is the first experimental study to show a direct correlation between storage duration and cardiac function. Infant and pediatric patients, and those receiving large volume and/or rapid transfusions may be sensitive to these effects.

## Introduction

More than 13 million whole blood and red blood cell units are transfused in the United States each year, with cardiac surgical procedures accounting for ∼20% of all blood transfusions(2, 10, 17, 33, 34, 51, 62). Many cardiac procedures mandate the use of blood and blood products in the preoperative, intraoperative and postoperative period, particularly with infant and pediatric patients for cardiopulmonary bypass circuitry priming(38, 62). Despite the frequency, transfusion of blood and blood products are not without risk(46, 58). Transfusion of red blood cells (RBC) in particular have been associated with increased morbidity and mortality, prolongation of hospital stay, and several different cardiac complications(30, 35, 36, 42, 44, 46, 52, 58, 59). Many investigators have suggested that RBC transfusion complications are due to the transfusion of RBCs close to their expiration (42 days), wherein the effects of the red cell storage lesion can contribute to the pathobiology of adverse reactions(7, 8, 14, 26, 40, 42, 44, 53, 54, 67). These pathobiological changes include clearance of storage-damaged RBCs, aberration of nitric oxide metabolism, trapping of RBCs by macrophages resulting in oxidative damage and impaired oxygen delivery, and an increase in circulating non-transferrin bound iron(29, 48, 53, 73). Briefly, over time, stored RBCs are depleted of ATP which alters the RBC cell membrane, resulting in hemolysis, the formation of red cell microvesicles, release of intracellular iron, decreased non-transferrin bound iron and the release of free hemoglobin. Further, the pH and electrolyte composition of the RBC unit also changes due to continued anerobic metabolism and dysfunction of cation transporters. The latter includes impairment of Na^+^/K^+^ ATPase(69), which leads to a progressive increase extracellular [K^+^] in the RBC unit supernatant(5, 28). Consequently, rapid or large volume transfusions of RBC units with elevated potassium levels can predispose patients to hyperkalemia, conduction abnormalities and cardiac arrest(7, 8, 24, 42, 54, 59). Although the incidence of transfusion-associated hyperkalemia is poorly defined and potentially underreported(42), Raza, et al. noted elevated K^+^ levels in >70% of adult trauma patients following transfusion(54), and Livingston, et al. observed hyperkalemia in 18-23% of pediatric trauma patients following transfusion(43). Transfusion-associated hyperkalemia resulting in cardiac arrest (TAHCA) is a recognized complication of massive transfusion in children, with a mean serum [K+] level of 9.2±1.8 mM in patients who experienced cardiac arrest(42). Some investigators suggest that the risk factors for TAHCA include the volume and rate of transfusion, storage age, and irradiation of RBCs – but the perceived risk and reason for such cardiac complications remains actively debated(4, 15, 28, 42).

Chronological storage age is one of the key factors that influences RBC quality and storage lesion severity(5, 12, 69). Despite this, blood banks often employ a “first-in, first-out” approach to reduce blood product waste and maintain an inventory supply to support emergency transfusions. Indeed, it is estimated that 10-20% of RBC units are transfused after 35-days of refrigerated storage, or near their 42-day expiration date(25). Some investigators have recommended a reduction in the maximum allowable storage time for RBCs due to quality concerns(29, 50, 53, 54, 61, 70, 71). Several clinical studies have raised concerns about the effects of the RBC storage lesion(8, 26, 37, 40, 42, 59, 75); however, the direct impact of RBC quality on cardiac health outcomes remains unclear. Identifying a mechanistic relationship between RBC quality and adverse cardiac endpoints has been hindered in the clinical setting by confounding factors, including disease diagnosis, age, rate/site of infusion, volume of transfusion per unit time, number of transfusions, bypass and cross-clamp time, secondary complications from surgery and concomitant medication administration. Recent randomized clinical trials have demonstrated that transfusion with fresh blood (1-10 days storage duration) does not decrease the risk of mortality compared with standard practice (2-3 weeks storage duration)(22, 27, 41, 63, 64). Although considerably less is known about the risk of transfusing RBCs near expiry (35-42 days), or the impact on secondary endpoints including cardiac complications(4, 39, 45, 55).

We aimed to address clinical concerns of bradycardia and cardiac arrest by investigating the direct relationship between RBC storage age and myocardial function using experimental models. We hypothesized that electrical conduction would be impaired in cardiac models exposed to the supernatant of ‘old’ RBC (sRBC) units close to expiration as compared with ‘fresh’ units, due in part to elevated extracellular potassium that can alter the myocardial resting membrane potential(3, 8, 21, 72). To test this hypothesis, electrophysiology parameters were measured using both an intact, isolated rat heart preparation and human stem-cell derived cardiomyocytes. Cardiac endpoints were measured at baseline, and again after exposure to sRBC collected from ‘fresh’ (day 7 post-donor collection), ‘old’ (day 30-40), or ‘expired’ units (day 50). We compared these results with those observed with hyperkalemia, a primary biomarker of RBC storage lesion severity(5, 12, 69).

M

## Materials and methods

### Red blood cell sample preparation

Red blood cell units (300 ± 50mL) from healthy donors were obtained from the American Red Cross or Children’s National Blood Donor Center. All blood units were O-negative, sickle-negative, non-irradiated, collected using standard single donor needle methods and stored in additive preservative solution (AS-1) according to standards of the American Academy Blood Banking requirements and the Food and Drug Administration(23). Single RBC units were aliquoted into small volume blood bags typically used for neonatal transfusion; each 100 mL aliquot was stored at 4-6^º^C in a research-grade, temperature monitored refrigerator according to standards(23). RBC units underwent gentle centrifugation (4^º^C, 20 min, 3700 rpm; Haemonetics) using accumulated centrifugal effect value of 6.5×10^7^ to separate and collect the supernatant (sRBC) 7-50 days post-donor collection; sRBC samples were used for subsequent experiments. Experiments were designed to study the impact of RBC storage lesion on cardiac electrophysiology, by comparing endpoints after exposure to ‘fresh’ sRBC (7 days post-donor collection), ‘old’ sRBC (30-40 days), or ‘expired’ sRBC (50 days).

### General protocol and biochemical analysis

Patients undergoing cardiac surgery or extracorporeal membrane oxygenation can receive large transfusion volumes equivalent to 60-70% of the patient’s total blood volume(19, 47). To mimic exposure, we estimated 10% supernatant volume exposure from reconstituted blood (1/2 volume packed RBCs [20-30% supernatant containing anticoagulant and 70-80% red blood cells] and 1/2 volume plasma). Accordingly, sRBC samples were diluted to 10% volume using Krebs-Henseleit buffered media (denoted in mM: 118 NaCl, 3.29 KCl, 1.2 MgSO_4_, 1.12 KH_2_PO_4_, 24

NaHCO_3_, 10 Glucose, 2 C_3_H_3_NaO_3_, 10 HEPES and 0.33 CaCl). Biochemical analyses were performed on each diluted sRBC sample, using an Epoc® point-of-care blood analysis system. Biochemical analyses were performed using a BGEM card (Seimens Diagnostics: SMNS10736382) to measure Na^+^, K^+^, Ca^2+^ and lactate levels.

### Intact, whole heart preparations

Animal protocols were approved by the Institutional Animal Care and Use Committee of the Children’s Research Institute, and followed the National Institutes of Health’s *Guide for the Care and Use of Laboratory Animals*.

Experiments were conducted using adult, female Sprague-Dawley rats (>8 weeks old, >200 g, Taconic Biosciences). Animals were housed in conventional rat cages in the Research Animal Facility under standard environmental conditions (12:12 hour light:dark cycle, 64 – 78F, 30-70% humidity, free access to reverse osmosis water, corn cob bedding and food (2918 rodent chow, Envigo). Animals were anesthetized with 3-5% isoflurane, the heart was excised and then transferred to a temperature-controlled (37°C), constant-pressure (70 mmHg) Langendorff-perfusion system for electrophysiology experiments (**Figure 1**). After isolating and transferring the heart to the perfusion system, excised hearts were perfused with Krebs-Henseleit buffer bubbled with carbogen (95% Oxygen, 5% CO_2_) throughout the duration of the experiment(31). Lead II electrocardiograms (ECG) were recorded continuously during sinus rhythm; ECG signals were analyzed to quantitate heart rate, atrioventricular conduction (PR interval), ventricular depolarization time (QRS width), ventricular repolarization (QTc) and arrhythmia incidence(32, 65). Biosignals were acquired in iox2 and ECG parameters were analyzed in ecgAUTO (emka Technologies).

**Figure 1.**
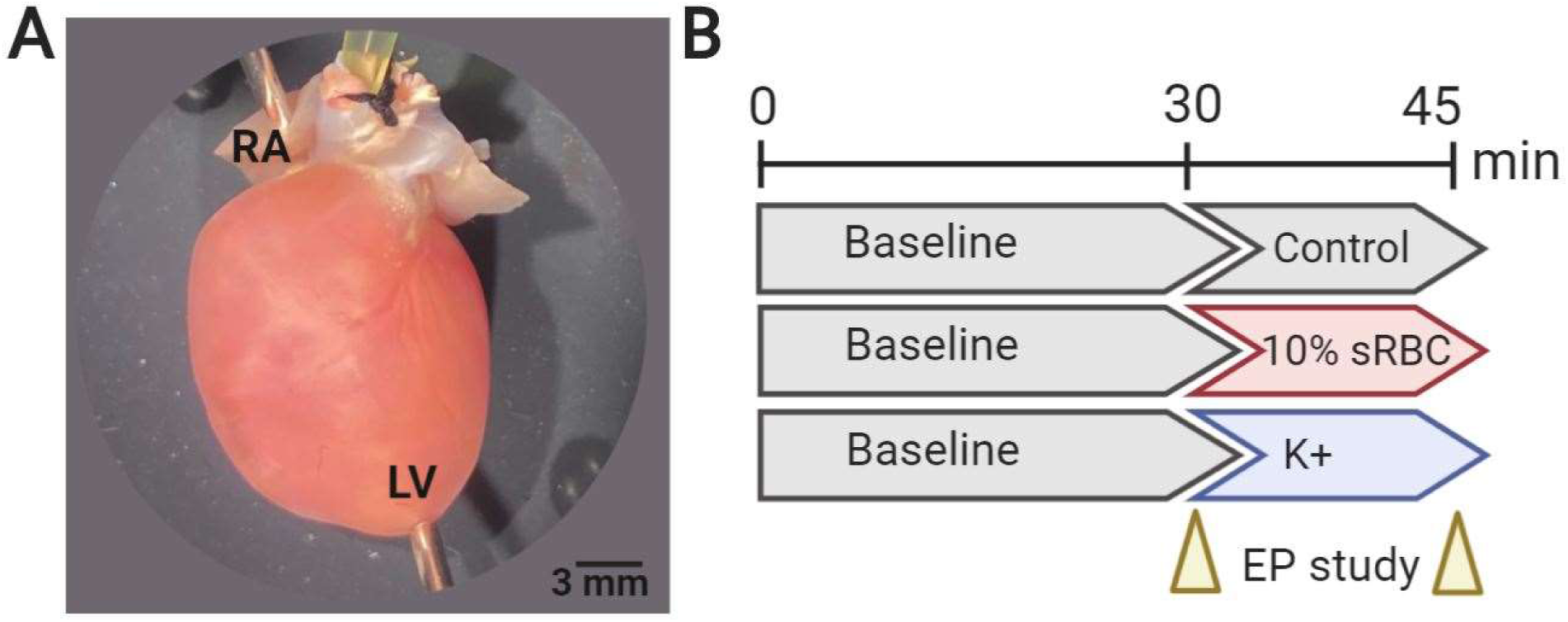
Heart preparation and experimental timeline. (A) Isolated, intact rodent heart with retrograde Langendorff-perfusion via an aortic cannula. Pacing electrodes were attached to the right atria (RA) and apex of the left ventricle (LV) to perform an electrophysiology study (EP). (B) Experimental timeline included 30-min perfusion with KH-media, containing 4.5 mM K^+^ (control), which commenced with an EP protocol. Thereafter, the media remained unchanged (control), supplemented with 10% sRBC, or supplemented with increasing potassium concentrations. The EP study was repeated again after 15-20 min, and results were compared to baseline.

### Electrophysiology measurements

To further investigate cardiac electrophysiology, a pacing protocol was implemented using stimulation electrodes positioned on the right atrium and the apex of the left ventricle (**Figure 1**)(32, 65, 66). A Bloom Classic electrophysiology stimulator (Fisher Medical) was set at a pacing current 1.5x the minimum pacing threshold (1-2 mA) with 1 msec monophasic pulse width. Sinus node recovery time (SNRT) was assessed by applying a pacing train of 150 ms (S1−S1) to the right atrium and measuring the time delay until the next spontaneous sinoatrial node-mediated activity. To determine the Wenckebach cycle length (WBCL), an S1-S1 pacing interval was applied to the right atrium; the pacing cycle length was decremented stepwise to pinpoint the shortest interval that resulted in 1:1 atrioventricular conduction. Next, an S1-S2 pacing interval was applied to the right atrium to determine the atrioventricular nodal effective refractory period (AVNERP). An S1-S2 pacing interval was applied to the left ventricle to find the shortest coupling interval that resulted in 1:1 ventricular depolarization, signifying the ventricular effective refractory period (VERP).

### Experimental timeline and treatment groups

Isolated, intact hearts were perfused with KH media for 30 min, followed by implementation of electrophysiology pacing protocols (‘baseline’). Hearts were then perfused for another 15-20 min, with either KH media alone (control), media supplemented with 10% sRBC (7-50 days post-donor collection), or media supplemented with elevated potassium concentrations (6-12 mM KCl). Electrophysiology protocols were performed a second time to determine the effects of sRBC treatment or hyperkalemia on electrical conduction (**Figure 1**). This protocol allowed each animal to serve as its own control, and account for experimental or animal variability.

### Human cardiomyocyte preparation and microelectrode array recordings

Human induced pluripotent stem cells differentiated into cardiomyocytes (hiPSC-CM; iCell cardiomyocytes) were plated onto fibronectin coated microelectrode arrays (Biocircuit MEA 24, Axion Biosystems), at a density of 30,000 cells per well. hiPSC-CM were maintained under standard cell culture conditions (37ºC, 5% CO_2_). hiPSC-CM formed a confluent contracting monolayer 2-4 days after plating (40-60 bpm) and MEA recordings were performed 7-10 days after plating to measure the spontaneous beating rate. hiPSC-CM were equilibrated in the MEA system for 15 min, and then the spontaneous beating rate was recorded (‘baseline’) using an integrated microelectrode array system (Maestro Edge, Axion) with temperature and gas control (37ºC, 5% CO_2_). Cardiomyocytes were then treated for 5 min with iCell maintenance media (control), media supplemented with 10% sRBC (7-40 days post-donor collection), or media supplemented with elevated potassium concentrations (9-12 mM). Spontaneous beating rate was also recorded 1 hr post-treatment and after washout. To account for cell plating variability, each treated cardiomyocyte monolayer was to baseline(11).

### Data analysis

Results are reported as mean ± standard error mean (n≥3 per group). Data normality was assessed by Shapiro-Wilk testing (GraphPad Prism). A two-tailed paired t-test was performed to compare endpoints before and after treatment, within the same heart (control media or sRBC). For hyperkalemia studies with multiple doses, statistical analysis was performed using either one-way analysis of variance or Kruskal-Wallis nonparametric test, with a false discovery rate (0.1) to correct for multiple comparisons. Significance was defined as *p<0.05.

## Results

### Storage age effects the biochemical composition of sRBC

The attributes of a stored blood product shifts as RBC quality declines, which can result in an accumulation of potassium in the supernatant(5, 12, 69). To measure the effect of storage time on the electrolyte composition of blood units, sRBC samples were collected from RBC units on day 7-50 post-donor collection, samples were diluted to 10% volume using pH-buffered KH media, and then electrolyte-gas measurements were performed on the diluted end product (**Figure 2**). Extracellular potassium levels were elevated in ‘old’ units as compared to ‘fresh’ units (day 7: 5.9±0.2 vs day 40: 9.7±0.4, p<0.0001); but, there was variability between age-matched units near expiry ranging from 8.5-11.9 mM [K^+^] in the 10% diluted end product (day 30-50). Lactate levels were also elevated in ‘old’ vs ‘fresh’ blood units (day 7: 0.8±0.1 vs day 40: 2.4±0.2 mM, p<0.0001).

**Figure 2.**
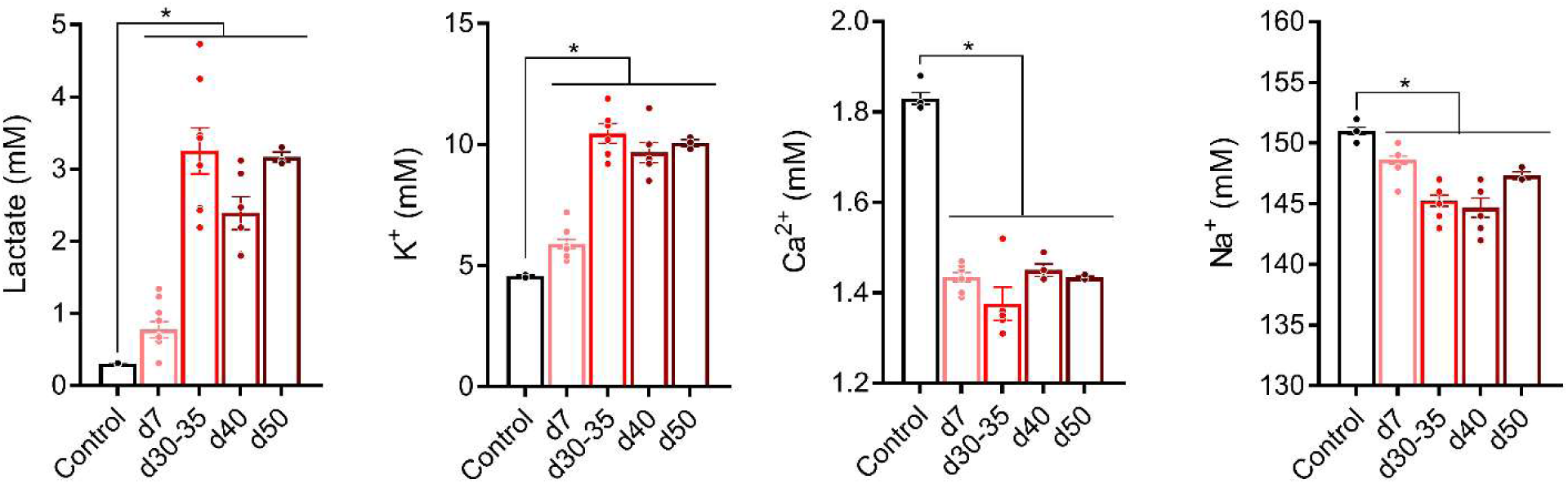
Biochemical composition of supernatant from red blood cell units (sRBC). Biochemical analyses of sRBC diluted to 10% volume in KH-buffered media. Storage age was associated with deviations in the electrolyte composition of sRBC samples. Mean ± SEM, *p < 0.05 relative to control (crystalloid KH perfusion buffer), n≥3 per time point.

### Storage age is associated with heart rate slowing and sinus node dysfunction

Cardiac complications from RBC transfusion include an increased risk of bradycardia and cardiac arrest(42, 54, 59, 67). These adverse outcomes may be precipitated by elevated extracellular potassium, which diminishes the myocardial resting membrane potential(21, 72). Accordingly, we assessed the impact of sRBC exposure on spontaneous heart rate and sinus node function in Langendorff-perfused hearts. Heart rate remained stable throughout the study when perfused with control media containing 4.5 mM K^+^ (baseline: 297±10 msec vs 45 min: 288±15 msec), and also remained stable when the perfusate was supplemented with 10% sRBC collected from RBC units aged 7-30 days (**Figure 3**). Similarly, sinus node function remained stable with control media perfusion (SNRT baseline: 223±14 vs 45 min: 238±9) and following perfusion with 10% sRBC collected from units aged 7-30 days (**Figure 3**). However, as RBC units neared expiration, sRBC exposure slowed the heart rate by 23% (baseline: 279±5 msec vs day 40: 216±18 msec, p<0.005). Additionally, sRBC from day 40 units had a significant effect on sinus node function, delaying the recovery time by 46% (SNRT baseline: 243±8 msec vs day 40: 354±23 msec, p<0.005). In the latter, the perfusate media had a mean potassium concentration near 10 mM (**Figure 2**). To measure the direct effect of hyperkalemia on automaticity and sinus function, a dose-response study was performed. As the potassium concentration increased from 4.5 to 12 mM, heart rate slowed (linear regression R^2^=0.92, p=0.01) and SNRT was prolonged (R^2^=0.86, p=0.02).

**Figure 3.**
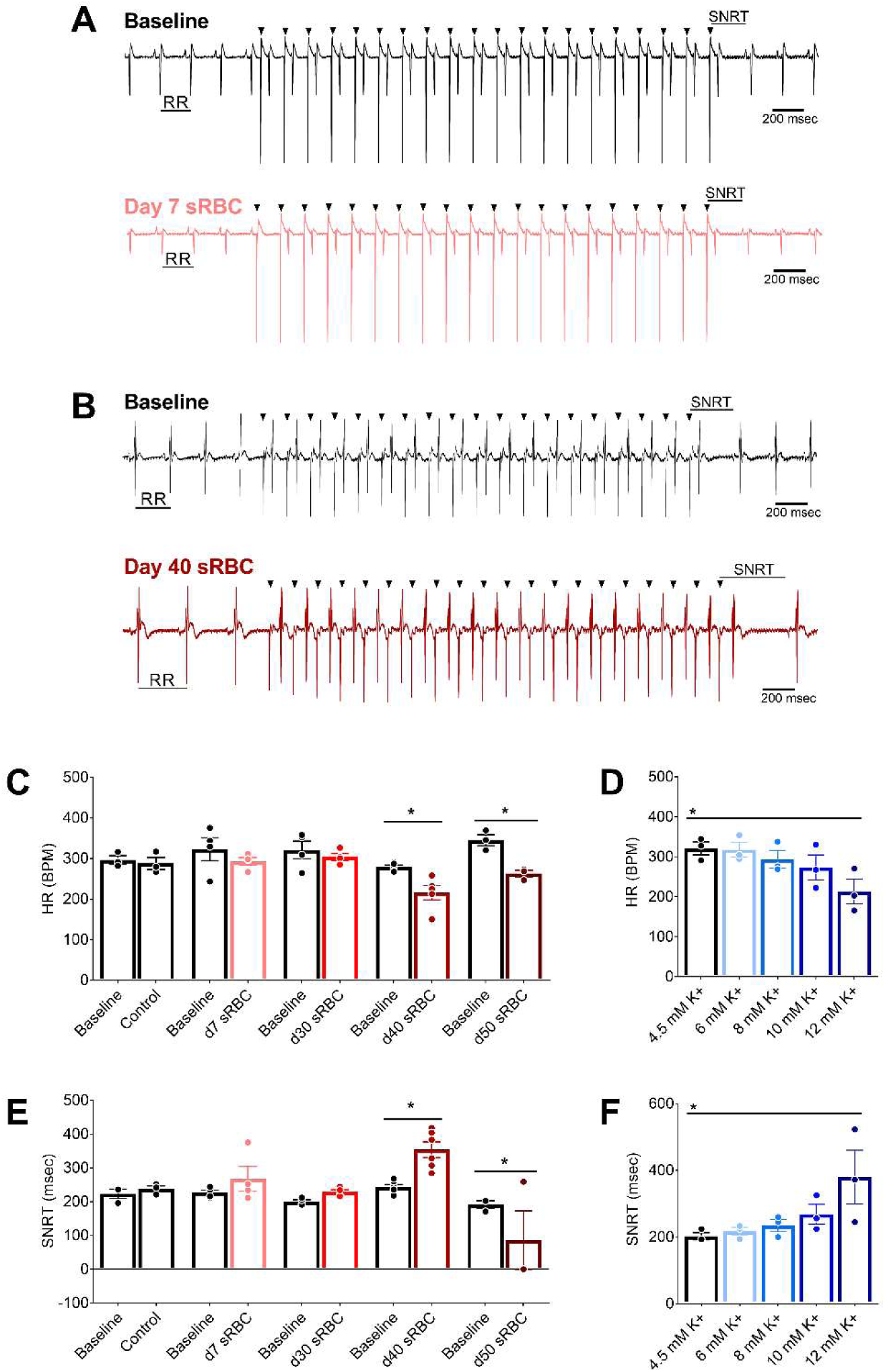
RBC storage age is associated with heart rate slowing and sinus node dysfunction. **(A)** Biosignals recorded from isolated hearts perfused with media supplemented with 10% sRBC collected from a day 7 unit, or **(B)** day 40 unit. Electrocardiograms were recorded during sinus rhythm (RR interval highlighted), followed by train of atrial paces (black arrows denote pacing spikes). Each atrial pace results in a ventricular response. Sinus node recovery time (SNRT) was measured from the last pacing spike to resumption of sinus rhythm. **(C)** Stable heart rate following exposure to RBC units aged 7-30 days, but bradycardia observed with sRBC collected from units aged ≥40 days. **(D)** Heart rate slowing observed at highest potassium concentration tested (12 mM K^+^). **(E)** Exposure to day 40 or 50 sRBC resulted in slowed sinus node recovery. **(F)** Increased SNRT also observed at highest potassium concentration tested (12 mM K^+^). Mean ± SEM, *p < 0.05.

### Storage age is associated with atrioventricular conduction slowing

Electrochemical gradients across the cardiomyocyte membrane are essential for cardiac excitation and electrical propagation. Atrial cardiomyocytes are particularly sensitive to deviations in these electrochemical gradients, and an increase in extracellular potassium can slow atrioventricular (AV) conduction(18, 21, 24). Atrioventricular conduction remained constant in hearts perfused with control KH media throughout the study (**Figure 4**), as determined by ECG parameters during sinus rhythm (PR time at baseline: 33±4 vs 45 min: 36±2). Similar results were observed before and after exposure to 10% sRBC samples collected from units aged 7-30, but significant slowing was observed after exposure to sRBC near or after expiration (PR time at baseline: 33±1 vs day 40: 41±3 msec, p<0.05; PR time at baseline: 37±1 vs day 50: 53±8 msec, p<0.005). AV node refractoriness was further interrogated by implementing an atrial pacing protocol to measure WBCL (S1-S1 pacing) and AVNERP (S1-S2 pacing). These parameters remained unchanged in hearts perfused with control media (WBCL baseline: 79±2 vs 45 min: 83±2; AVNERP baseline: 64±5 vs 45 min: 67±4) and hearts exposed to sRBC from ‘fresh’ 7-day units (**Figure 5**,**6**). Exposure to day 30 sRBC resulted in a modest increase in AV node refractoriness, increasing WBCL by 9%. Effects on the AV node were more pronounced after exposure to day 40 sRBC which increased AVNERP by 21% (baseline: 77±2 vs day 40: 93±7 msec, p=0.01) and WBCL by 19% (baseline: 90±1 vs day 40: 107±3msec, p<0.001). These effects were further exacerbated in units stored past expiration (78% increase in WBCL and 66% increase in AVNERP, baseline vs day 50 sRBC; **Figure 5**,**6**).

**Figure 4.**
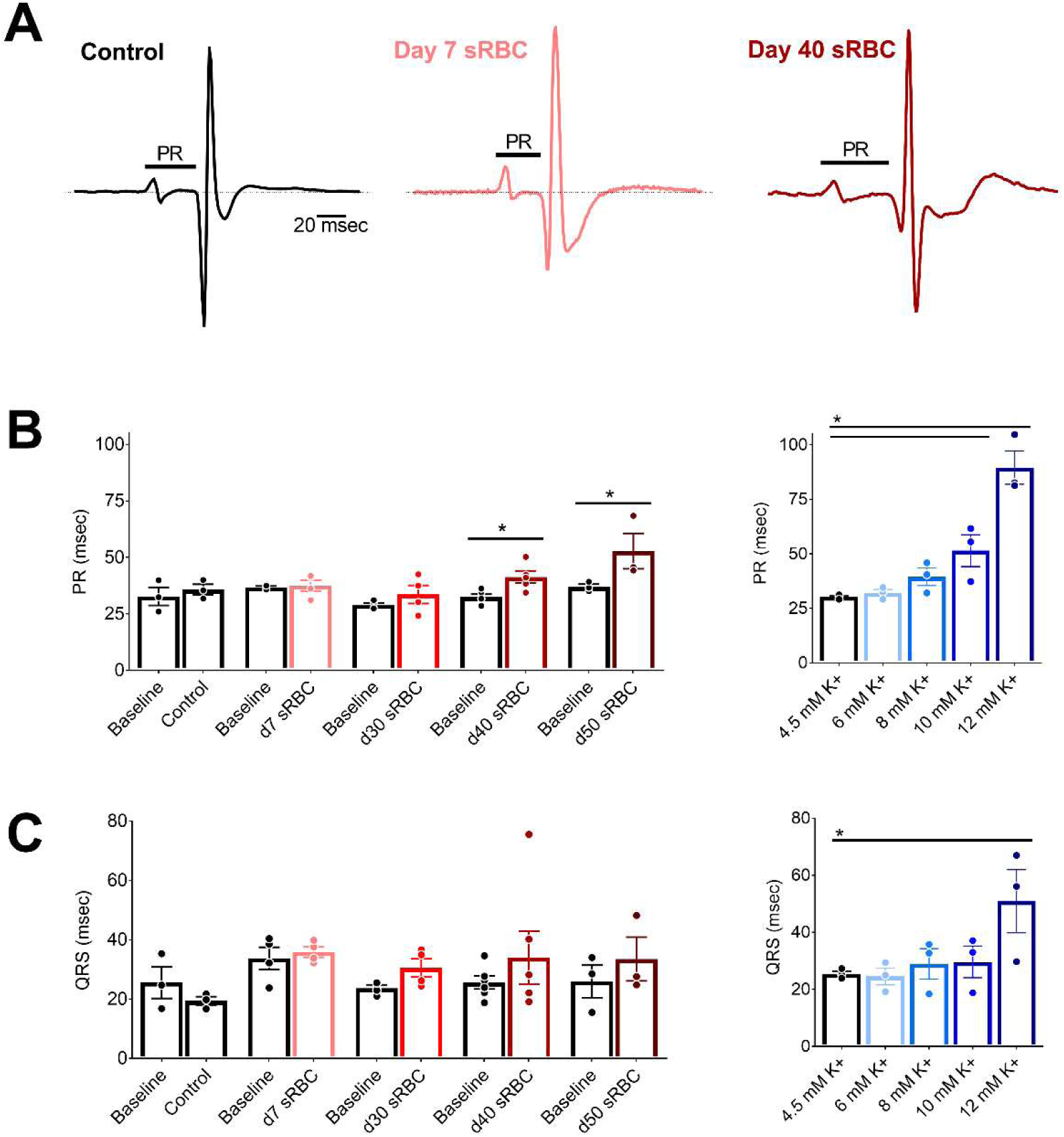
RBC storage age is associated with slowed atrioventricular conduction. **(A)** Electrocardiograms recorded during sinus rhythm from isolated hearts perfused with control media (left), media supplemented with 10% sRBC collected from a day 7 unit (middle) or day 40 unit (right). PR interval time is denoted. **(B)** Atrioventricular conduction slows in the presence of day 40 and day 50 sRBC, or 10-12 mM K^+^. **(C)** Exposure to sRBC units had no measurable effect on ventricular depolarization time (QRS) during sinus rhythm. Mean ± SEM, *p < 0.05.

**Figure 5.**
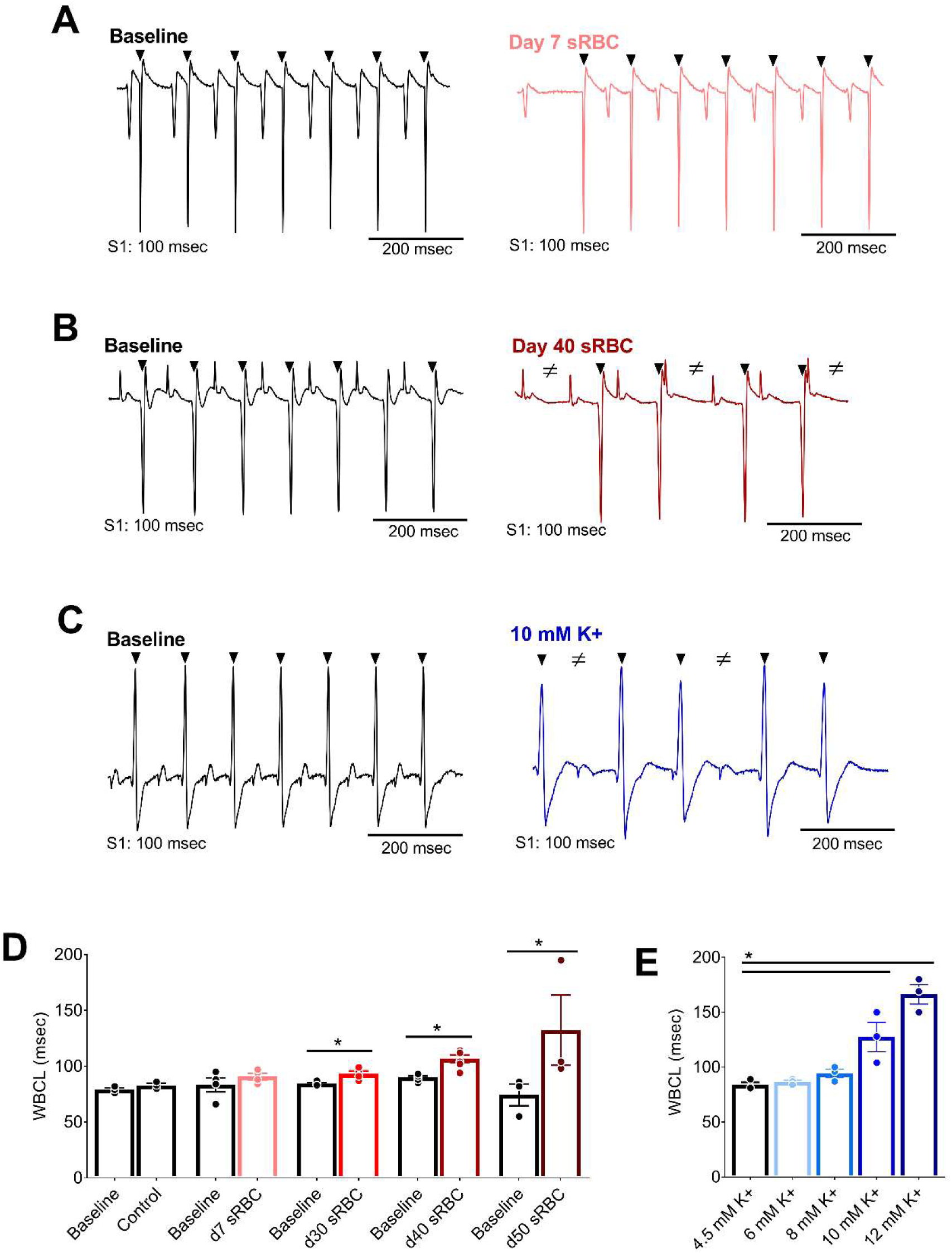
RBC storage age is associated with increased refractoriness of the AV node. **(A)** Biosignals recorded with atrial pacing (S1-S1) to measure Wenckebach cycle length (WBCL) in isolated hearts in the presence of day 7 sRBC, **(B)** day 40 sRBC, or **(C)** 10 mM K^+^. **(D)** Slowed atrioventricular node conduction following exposure to sRBC from units 30-50 days old, but not ‘fresh’ day 7 units. **(E)** Slowed atrioventricular conduction following exposure to 10-12 mM K^+^. Arrows denote ventricular response to atrial pacing at S1 (black) pacing cycle length. ≠ denotes failed conduction. Mean ± SEM, *p < 0.05.

**Figure 6.**
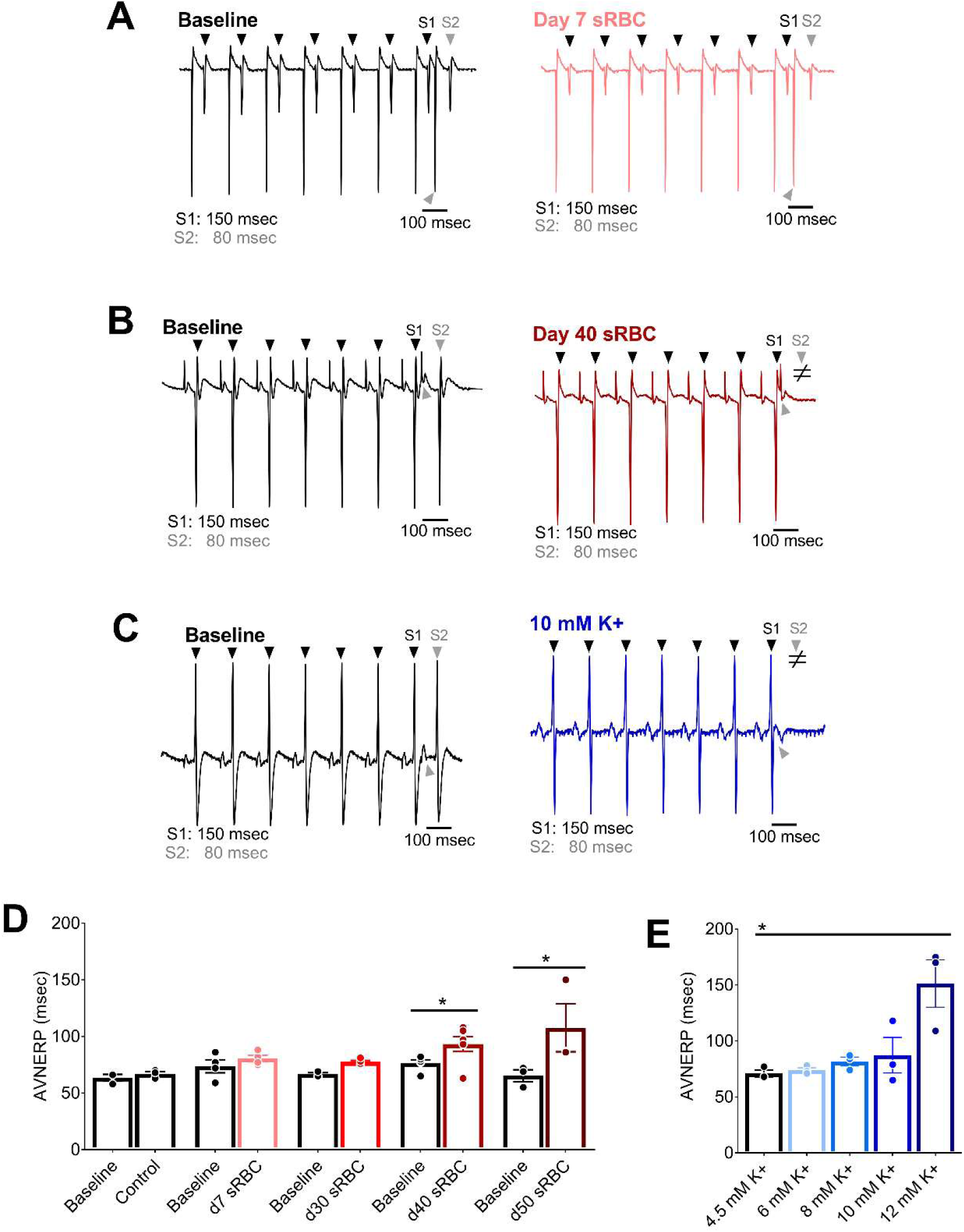
RBC storage age is associated with an increased AV node effective refractory period. **(A)** Biosignals recorded with atrial pacing (S1-S2) to pinpoint atrioventricular node effective refractory period (AVNERP) in the presence of day 7 sRBC, **(B)** day 40 sRBC, or **(C)** 10 mM K^+^. **(D)** AVNERP did not change after exposure to day 7-30 sRBC, but increased with day 40 and day 50 sRBC exposure. **(E)** AVNERP increased with severe hyperkalemia. Arrows denote ventricular response to atrial pacing at S1 (black) or S2 (gray) pacing cycle length. ≠ denotes failed conduction. Mean ± SEM, *p < 0.05.

As anticipated, a dose response relationship was observed when the potassium concentration was increased in the perfusate media, resulting in prolonged atrioventricular conduction time and increased AV node refractoriness. As the potassium concentration increased from 4.5 to 12 mM, a progressive increase in PR duration (R^2^=0.85, p<0.05) was observed (**Figure 4**). At 10 mM K^+^ (a concentration comparable to day 40 sRBC-supplemented media), a 51% increase in WBCL was observed (4.5 mM: 84±3 to 10mM: 127±13 msec, p<0.005), but changes in AVNERP were only observed at 12 mM K^+^ (4.5 mM: 71±3 to 12 mM: 151±21 msec, p<0.005; **Figure 5**,**6**). The latter suggests that other factors or substances in the RBC supernatant may also contribute to conduction slowing.

### Storage age increases ventricular refractoriness

Severe hyperkalemia is associated with decreased sodium channel availability and slowed conduction velocity, which results in QRS widening and may precipitate ventricular tachyarrhythmias(18, 21, 24). In our study model, exposure to sRBC-supplemented media did not significantly prolong the QRS duration (baseline: 26±2 msec vs day 40: 34±9 msec; **Figure 4**), QTc duration (baseline: 169±9 vs day 40: 172±11 msec) or increase the incidence of ventricular tachyarrhythmias (data not shown). Further, we were not able to establish a trend toward QRS prolongation with increasing potassium concentration (R^2^=0.72, p=0.07), QTc duration (R^2^=0.67, p=0.67) or an increased incidence of ventricular tachyarrhythmias – which may be attributed to limitations in our model system. Indeed, ventricular activation and early repolarization can occur simultaneously in the rodent heart – which can influence the QRS complex and result in indistinct T-waves(6). Moreover, the rodent myocardium is less than ideal for assessing arrhythmia incidence due to its small size and resiliency to fibrillation(6). As another indicator of ventricular repolarization time, we implemented a pacing protocol to pinpoint ventricular refractoriness. A marginal increase in extracellular potassium can hasten repolarization and shorten action potential duration time – but severe hyperkalemia increases potassium channel conductance, lengthens action potential duration, and increases ventricular refractoriness(49, 72). As expected, control media perfusion resulted in stable VERP measurements throughout the study (VERP baseline: 45±5 vs 45 min: 46±2 msec). VERP measurements were unchanged in heart preparations exposed to sRBC from day 7-30 RBC units (**Figure 7**), but VERP increased by 96% following exposure to day 40 sRBC (baseline: 50±3 vs day 40: 98±10 msec, p<0.0001) and 145% after exposure to expired units (baseline: 51±8 vs day 50: 126±25 msec, p<0.0001). This increase in ventricular refractoriness may be explained, at least partly, by the increase in extracellular potassium levels. In dose response studies, increasing potassium concentration (4.5 to 12 mM) also resulted in a progressive increase in VERP (linear regression, R^2^=0.91, p=0.01).

**Figure 7:**
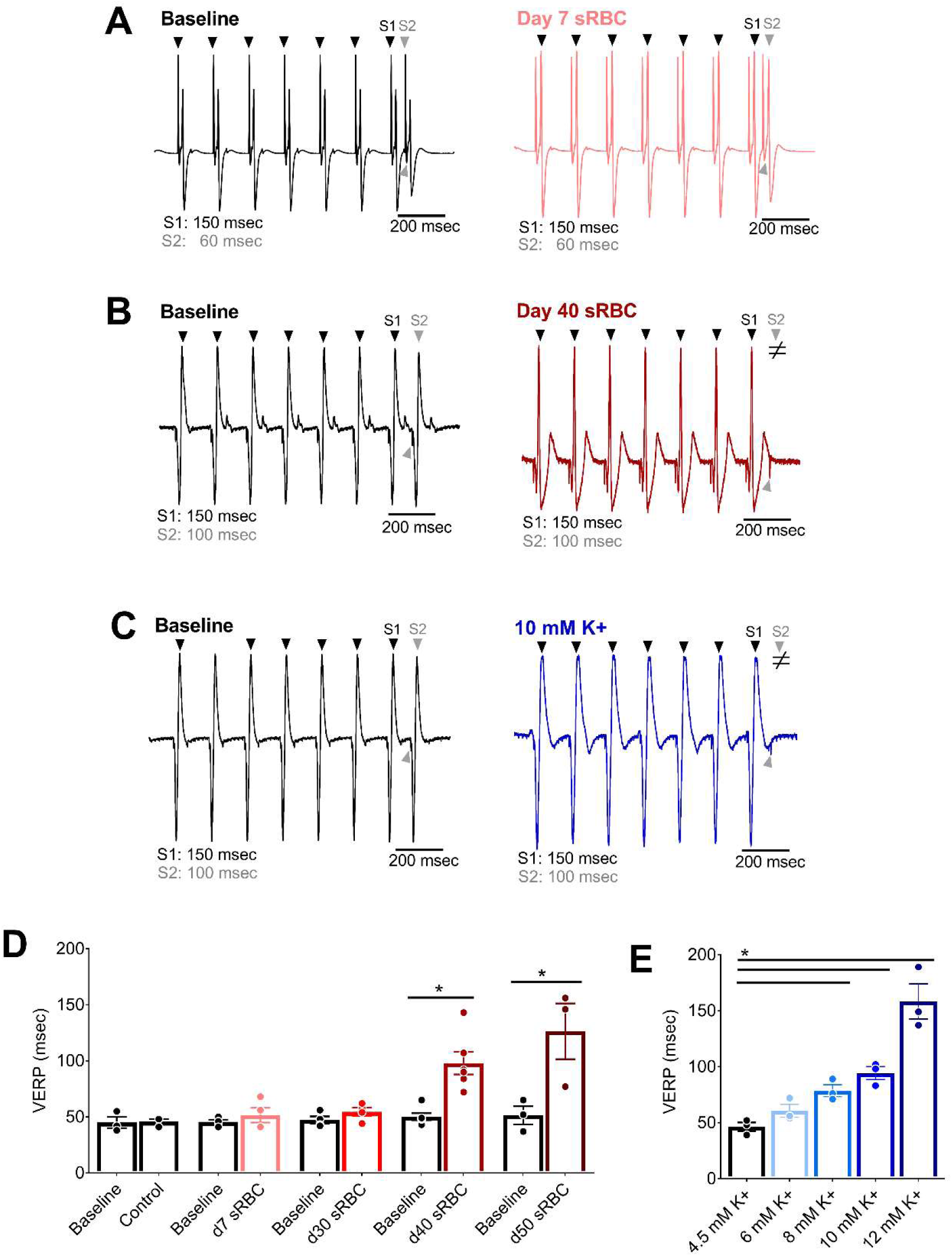
RBC storage age is associated with increased ventricular refractoriness. **(A)** Biosignals recorded with ventricular pacing (S1-S2) to pinpoint the ventricular effective refractory period (VERP) in isolated hearts perfused with media supplemented with 10% sRBC collected from a day 7 unit, **(B)** day 40 unit, or **(C)** 10 mM K^+^. **(D)** Ventricular refractoriness was unchanged after exposure to day 7-30, but increased with day 40-50 sRBC and **(E)** media supplemented with 8-12 mM K^+^. Arrows denote ventricular response to pacing at S1 (black) or S2 (gray) pacing cycle length. ≠ denotes failed conduction. Mean ± SD, *p < 0.05.

### Human cardiomyocytes are susceptible to electrical disturbances

Rodent models are frequently employed in cardiovascular research studies, although species-specific differences in ion channel expression are established(20, 74). Accordingly, we performed a follow-up study using human cardiomyocytes (hiPSC-CM) to validate the effects of sRBC exposure. Using a microelectrode array (MEA) system, we noted an increase in the beating rate of hiPSC-CM over time when treated with day 7 sRBC (5min: 12±6% rate increase p=0.09 vs 60min: 33±5% p<0.005, **Figure 8**). Conversely, cardiomyocytes demonstrated bradycardia after exposure to ‘older’ sRBC products, which was more severe than reported in the whole heart experiments. The spontaneous beating rate of hiPSC-CM decreased by 47±7% in day 35 samples and 75±9% in day 40 samples relative to baseline measurements (p<0.0001). Significant slowing in the spontaneous beating rate was also observed with increasing potassium concentrations (4.5-12 mM K^+^; R^2^=0.999, p=0.01). Notably, treatment did not appear to have a lasting effect on cardiomyocyte viability, as the beating rate quickly returned to normal after washing out the sRBC or hyperkalemic media (**Figure 8**).

**Figure 8.**
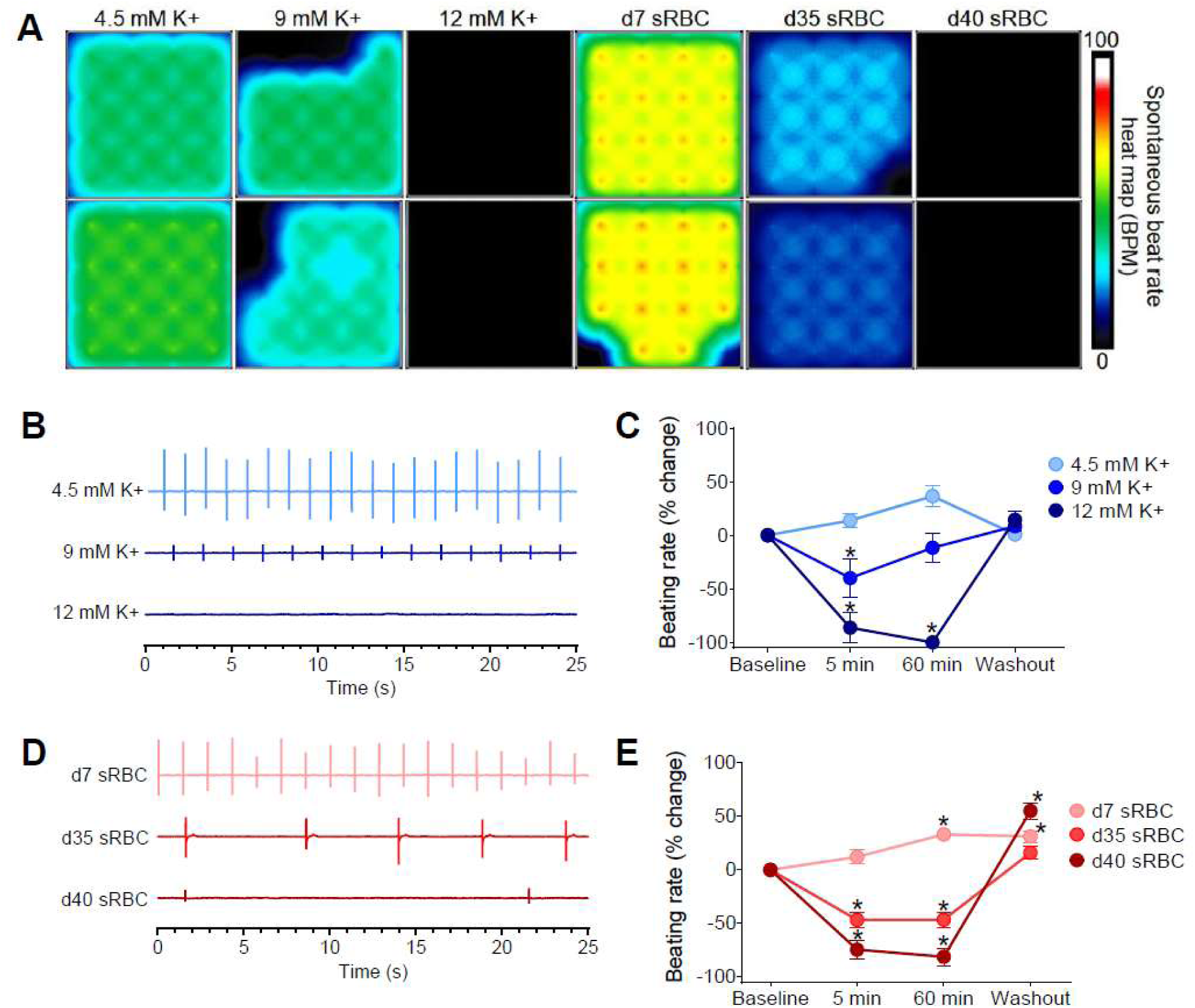
Reduced automaticity in human cardiomyocytes. **(A)** Microelectrode array heat map shows 16-electrode recordings from cardiomyocytes treated with control media (4.5 mM K^+^), media with increasing potassium concentrations (9-12 mM K^+^) or 10% sRBC collected from RBC units aged 7-40 days. The heat map corresponds to the spontaneous beating rate. **(B)** Biosignals recorded from human cardiomyocytes show a decline in beating rate with elevated potassium concentrations. **(C)** Percent change in beating rate following treatment with elevated potassium concentrations, compared to baseline. **(D)** Biosignals show a decline in the beating rate with ‘older’ sRBC samples (day 35-40) but not ‘fresh’ sRBC samples (day 7). **(E)** Percent change in beating rate following sRBC treatment, compared with baseline. Mean ± SEM, n≥12, *Significantly different from baseline, p < 0.05.

## Discussion

Clinical case reports have documented transfusion-associated hyperkalemia, which can lead to conduction disturbances, ventricular tachycardias, and/or cardiac arrest(3, 7, 8, 24, 42, 54, 59). Further, studies suggest that transfusion-associated adverse events may be associated with the storage age of blood products, as RBCs undergo a cascade of morphological, biochemical and metabolic changes over time that are collectively termed the ‘RBC storage lesion’ or ‘metabolic aging’(7, 42, 54, 60). This study is the first to demonstrate that ‘older’ blood products may directly impact myocardial automaticity and electrical conduction, using experimental cardiac models. Importantly, we show that supernatant collected from ‘fresh’ RBC units (7 days post-donor collection) had no effect on heart rate, sinus node function, atrial or atrioventricular conduction, or myocardial refractoriness in an isolated, whole heart model. A follow-up study in human cardiomyocytes revealed that supplementation with 10% sRBC from ‘fresh’ units (day 7) had a modest increased the spontaneous beating rate over time, which may be attributed to mild hyperkalemia (6.0±0.6 mM K^+^). In comparison, whole heart preparations exposed to supernatant from aged RBC units (>30 days post-collection) displayed bradycardia, slowed atrial and atrioventricular conduction, and an increase in the refractoriness of the ventricle and AV node. Notably, other groups have suggested that the maximal allowable red cell storage duration be reduced from 42 to 35 days, due to increased hemolysis and a sharp increase in nontransferrin-bound iron after 5 weeks in refrigerated storage(53). Although we did not measure either free iron or non-transferrin bound iron levels in this study, our results closely align with this conclusion, as electrophysiological disturbances were predominately observed in units stored 30+ days post-donor collection.

### Mechanistic links between RBC transfusion and adverse cardiac outcomes

Blood transfusion complications include an increased risk of bradycardia and cardiac arrest, which may be precipitated by an elevated potassium level in the supernatant of RBC units(3, 8, 42, 59, 67). As extracellular potassium increases, electrochemical gradients are diminished and the cardiomyocyte resting membrane potential becomes less negative(18, 49, 72). Accordingly, mild hyperkalemia can enhance cardiomyocyte excitability – similar to our observation with day 7 sRBC treatment in human cardiomyocytes. But, with more severe hyperkalemia, the change in resting membrane potential decreases the availability of voltage-gated sodium channels that are critical to depolarization and myocardial excitability(72). Accordingly, severe hyperkalemia is marked by sinus node dysfunction and sinus arrest(21). Similar observations were observed in our study when cardiac preparations were exposed to increasing potassium concentrations, a prominent biomarker of red cell storage lesion that can, at least in part, contribute to the electrical disturbances observed in this study.

As described above, hyperkalemia shifts the resting membrane potential and reduces the availability of voltage-gated sodium channels. As the action potential upstroke slows, electrical conduction slows, which manifests as a prolongation of P-waves, PR interval and QRS interval time(18, 49, 72). Atrial cardiomyocytes are the most sensitive to elevated potassium concentrations – followed by the ventricular myocardium and then specialized conductive tissue, including the sinoatrial node and bundle of His(18, 49, 72). Accordingly, electrical disturbances attributed to high [K^+^] are initially observed as widened p-waves with shorter amplitudes, followed by atrioventricular and ventricular conduction delays as extracellular [K^+^] continues to increase. Instead of a gradual change in cardiac parameters, we observed a global depression in electrical conduction that was largely limited to sRBC samples near expiration and/or 10-12 mM K^+^ perfusion. The latter may be attributed to the sensitivity of our model system(6), species-specific differences in ion channel expression and electrophysiology(20, 74), and/or other attributes of the RBC storage lesion (e.g, lactate, free-iron, plasticizer leaching) that may have additional effects on cardiac electrophysiology(13, 14, 29, 32, 53).

Although not investigated in the present study, phthalate chemical exposure is another potential contributor to heart rate slowing and sinus node dysfunction. Phthalate chemicals are frequently used as plasticizers in blood bags, and studies have shown that storage age is associated with an accumulation of harmful phthalate chemicals in the supernatant of stored RBC products (18-fold increase, day 5 vs 42 post-donor collection)(13). Phthalate chemical exposure has been associated with bradycardia in *in vivo*(56), *in vitro*(57) and using an isolated heart model(1). Moreover, our laboratory previously reported that phthalate plasticizers can lead to sinus node dysfunction in an isolated heart model, delaying SNRT by 54% compared with control(32). Additional studies are needed to investigate the additive effects that may result from hyperkalemia and phthalate chemical exposure.

### Clinical Implications

In the current study, we focused our attention on hyperkalemia as a plausible mechanism for the electrophysiology disturbances observed in our model system after exposure to ‘old’ RBC samples. Hyperkalemia has been reported in >70% of adult trauma patients following transfusion(54), and observed in 18-23% of pediatric trauma patients following transfusion(43). Moreover, Smith, et al. reported that an increase in serum potassium levels (5.9-9.2 mEq/l) was associated with a higher risk of cardiac arrest(59), which is more likely to occur following rapid transfusion, large volume transfusion, or in cases of low cardiac output that impairs the redistribution of potassium(7, 42). Potential solutions to help mitigate the risk of hyperkalemia include prebypass filtering(16), washing RBCs(67) or limiting RBC storage duration(40, 42, 53, 54, 59). Notably, longer blood storage duration has been associated with suboptimal outcomes in high-risk pediatric surgery cases(44) and cardiac operations(40, 52). Recent randomized controlled trials have indicated that transfusion of ‘fresh’ blood (e.g., 1-10 days) does not decrease the risk of mortality when compared to standard of care (e.g., 2-3 weeks)(22, 27, 41, 63, 64). However, much less is known about the safety of prolonged RBC storage (e.g., 30-42 days) or the impact of ‘old’ blood products on secondary cardiac endpoints(4, 55). Accordingly, expert panels have highlighted the lack of evidence-based data to reach consensus on the safety of RBC storage age in relation to critically ill children, including those undergoing surgical repair for congenital heart defects or those undergoing extracorporeal membrane oxygenation(9, 68). The presented study highlights the importance of studying the direct impact of RBC storage lesion on end-organ function, with an emphasis on cardiac electrophysiology given the sensitivity of the heart to electrolyte disturbances.

## Limitations

The scope of our study was limited to the effects of acute cardiac exposure to supernatant collected from RBC units. Whole heart and cardiomyocyte models were used to investigate the direct effects of sRBC-mediated biochemical disturbances on electrical activity. However, *in vitro* and *ex vivo* results may differ from those observed *in vivo*, with an intact vascular and autonomic nervous system. To mimic patient exposure following a large transfusion, we estimated 10% supernatant volume exposure from reconstituted blood – based on volumes reported in cardiac surgery and/or extracorporeal membrane oxygenation studies.

Additional studies are warranted to assess additional effects that may result from reconstituted blood containing aged RBCs, or the risk to sensitive populations including those with low cardiac output.

## Acknowledgements

The authors gratefully acknowledge Dr. Luther Swift for experimental technical assistance, Drs. Nobuyuki Ishibashi and Takuya Meada for assistance with electrolyte measurements, and Drs. Pranava Sinha and Charles Berul for helpful discussions. We also acknowledge Dr. Meghan Delaney and Antoine Tavares Da Souza in the Children’s National Blood Bank for their assistance with procuring, storing, and collecting RBC supernatant products for this study.

## Funding Acknowledgements

This work was supported by the National Institutes of Health (R01HL139472 to NGP), Sheikh Zayed Institute for Pediatric Surgical Innovation, and the Children’s National Heart Institute. This publication was also supported by the Gloria and Steven Seelig family.

